# Habituation to visual stimuli is personality-independent in a jumping spider

**DOI:** 10.1101/2023.05.09.539210

**Authors:** Narmin Ilgar Beydizada, Francesco Cannone, Stano Pekár, David Baracchi, Massimo De Agrò

**Affiliations:** Department of Botany & Zoology, Faculty of Science, Masaryk University, Kotlářská 2, 611 37 Brno, Czech Republic; Department of Biology, University of Florence, Via Madonna del Piano, 6, 50019 Sesto Fiorentino, Italy

**Keywords:** *Menemerus semilimbatus*, boldness, visual detection, response rate, dis-habituation, habituation

## Abstract

Jumping spiders display some of the richest visually-mediated behaviors in nature. Vision is indeed the most important sensory modality in these spiders where motion detection and response to visual stimuli allow key behaviors such as hunting, escaping from predators, and mating. These spiders have been used in various experiments demonstrating the existence of good associative learning and memory abilities, whose mechanism parallels that found in vertebrates. Here we focused on the habituation and dis-habituation (H/DH) paradigm, indicating either a gradual decrease in responsiveness to repeated visual stimuli (H), or a recovery of the habituated stimulus (DH). H is an elementary form of non-associative learning and memory, which is expected to vary from individual to individual. The link between personality and H/DH has been shown in many vertebrates, but rarely in invertebrates. To tackle this question we tested whether personality affects H/DH in the jumping spider *Menemerus semilimbatus.* In our protocol, habituation was assessed by presenting repeatedly a visual stimulus on a screen to spiders tethered on a locomotor compensator. In the same individuals, personality (namely boldness) was assessed in a walking arena equipped with a shelter. We found that *M. semilimbatus* habituated and dishabituated to our visual stimulus and that they differed in personality along a shy-bold axis. However, contrary to our expectations, personality was not related to learning. We discussed the results and speculated that the nature (neutral value) of the stimulus might have played a role in making learning independent from personality.

## 2 Introduction

For animals, it is crucial to be able to detect and memorize important stimuli, and thus to shift and/or improve their behavioral strategies as a result of experience and to adapt to constantly changing circumstances. However, memorizing every single object or event would not only be impossible but would require too many resources and time without any ultimate benefit for the individual. Therefore, given the limited storage capacity and computing power of brains, animals necessarily filter information, discerning relevant from irrelevant ones (Shettleworth, 1993).

The ability to learn and memorize is ubiquitous in nature and fundamental to any taxa, from vertebrates to invertebrates and plants (Gagliano, 2018; Menzel & Benjamin, 2013; Purves et al., 2019; Squire & Kandel, 2003). One of the most elemental forms of learning is habituation (Giles & Rankin, 2009), which refers to the reduction in physiological responses to a repeated homotypic (same) stimulus. Habituation is reversible: if the stimulus is withheld, the response recovers over time (spontaneous recovery). As a historical example, in *Aplysia*, repeated electric shocks or tactile stimulations that initially generated a defensive withdrawal reflex induced habituation and response extinction (Rankin & Carew, 1987). After the stimulus became habituated, the re-presentation of the same stimulus (electric shock or physical touch) after some time evoked (reactivated) the same defensive behavior again, i.e., induced dis-habituation.

Central to the present study, habituation can also be induced by the repeated presentation of stimuli with no intrinsic value. In this case, habituation attests to the animal’s ability to learn that a particular stimulus is indeed irrelevant (Thompson & Spencer, 1966) (i.e., when the animal realizes the stimulus has no effect or value, the responses drop). After habituation has occurred, dis-habituation can be induced by presenting a novel stimulus (Giles & Rankin, 2009). To be considered novel, a stimulus may vary from the habituation one even by a single characteristic (shape, size, intensity, motion direction, speed, etc.). In this case, the occurrence of dis-habituation is an indication of an animal’s ability to discriminate between the two stimuli.

Individuals of the same species may vary consistently in the way they perceive, recognize/discriminate objects or stimuli (Augustsson & Meyerson, 2004; Carere & Locurto, 2011; Guillette et al., 2015; Matzel et al., 2011). These consistent inter-individual differences can be referred to as a personality trait – namely average-level stable, long-term repeatable behavioral, emotional, and physiological differences in suites of traits across time and/or contexts among individuals of the same species (Réale et al., 2007). Personality traits such as boldness, aggressiveness, and activity level (Bell et al., 2009; Carter et al., 2013) have been reported to have a profound effect on many aspects of animals’ behavior and cognition (Dougherty & Guillette, 2018; Griffin et al., 2015; Matzel et al., 2011), including associative and non-associative learning abilities (Carere & Locurto, 2011; Allan et al., 2020, Finke et al., 2021, 2023). A recent meta-analysis performed on 19 species across a wide spectrum of taxa, including insects, showed that bold animals are generally faster learners (Dougherty & Guillette, 2018). The link between personality differences and habituation has been mainly documented in vertebrate taxa (e.g., in mammals: Mazza et al., 2018; fish: Etheredge et al., 2018; birds: Dissegna et al., 2022; Jha & Kumar, 2017; humans: LaRowe et al., 2006; O’Gorman, 1977). However, only a very limited number of invertebrate models are represented in the study of habituation itself (Baglan et al., 2017; Pietrantuono et al., 2021; Engel & Wu, 2009; Haupt, 2004; Haupt & Klemt, 2005; Pinsker et al., 1970; Plowright et al., 2006, Baracchi et al., 2018), and effect of individual differences (personality) in habituation and dis-habituation on invertebrate models remains unexplored.

Spiders are one of the most behaviorally diverse and successful groups of terrestrial predators (Coddington & Levi, 1991) which challenges the conventional assumption of cognitive limitations in small animals (Cross et al., 2020; Jakob et al., 2011). The presence of personality types and their link with various behaviors has been observed in many spider species (Beydizada & Pekár, 2023; Chang et al., 2018, 2017; Foellmer & Khadka, 2013; Grinsted & Bacon, 2014; Liedtke et al., 2015; Michalko et al., 2017; Royauté et al., 2015). Among spiders, salticids display some of the richest behaviors and varied personality types (Chang et al., 2018, 2017; Chia-Chen, 2019; Harland & Jackson, 2004; Liedtke et al., 2015). As active hunters, salticids must selectively attend to a wide range of visual stimuli. Their visual system is impressive (Harland et al., 2012), divided into functionally distinct eye pairs. Some of these are specialized in motion detection and recognition (De Agrò et al., 2021; Spano et al., 2012; Zurek et al., 2010; Zurek & Nelson, 2012a) while others are equipped with long eye tubes, capable of saccadic tracking (Bruce et al., 2021; Jakob et al., 2018; Land, 1969a, 1969b), and are specialized in object recognition (Bednarski et al., 2012; Dolev & Nelson, 2016, 2014; Rößler et al., 2022). Jumping spiders have been tested in various experiments demonstrating the presence of associative learning and memory (Cross & Jackson, 2017; De Agrò, 2020; De Agrò et al., 2017; Hoefler & Jakob, 2006; Jakob & Long, 2016; Peckmezian & Taylor, 2015; Skow & Jakob, 2006; Taylor et al., 2016). Response decrement over time has been shown previously in some jumping spiders (De Agrò et al., 2021; Humphrey et al., 2018, 2019; Nelson et al., 2019; Zurek et al., 2010; Zurek & Nelson, 2012a). Habituation instead has been directly tested only in a salticid, *Trite planiceps* (Melrose, 2015; Melrose et al., 2018).

In this study, we investigated whether a personality trait (i.e., boldness) is related to the habituation and dis-habituation rates to visual stimuli in the jumping spider *Menemerus semilimbatus*, Hahn (Salticidae). We hypothesized that bolder individuals would have a higher probability of responding to any stimulus (i.e., those who initially react more likely) and maintain their response rate higher for a longer time compared to shy individuals.

## 3 Material and Methods

### 3.1 Study organism

We collected 91 jumping spiders of the species *Menemerus semilimbatus* (mean ± SEM of prosoma length = 2.852 ± 0.639 mm) of two developmental stages (70 juveniles, 10 adult males, and 11 adult females) in the autumn (September-October) 2022 around the University campus. Spiders were transferred to the laboratory and measured with millimeter graph paper (prosoma length and width in mm). They were then housed singly in clean boxes (75 × 65 × 155 mm). Shelter and water for each spider were provided. Spiders were kept under an automatized light regime (LD=12:12).

### 3.2 Experimental procedure

Each spider was kept in the lab in total for five days, during which it underwent a visual habituation/dis-habituation (H/DH) test and two personality assays.

#### 3.2.1 Day 1: Subject capture and preparation

A spider was captured and immediately prepared for the subsequent test by fixing a magnet to its carapace. To do so, a spider was first gently constrained between a sponge and a latex film. The latter presented a hole in correspondence with the location of the spider’s head. Here, a 1×1mm cylindrical neodymium magnet was glued using a UV-activated resin. .

#### 3.2.2 Day 2: Habituation/Dis-habituation (H/DH) test

The spiders were subjected to a visual H/DH test. This was performed using a spherical treadmill identical to the one described in De Agrò et al. (2021). In brief, each spider was attached to the end effector of a micro-manipulator by means of the magnet glued to their cephalothorax. Then, the spider was positioned on top of a styrene ball, made frictionless thanks to the continuous flow of compressed air. The distance between the spider and the ball was set to mimic the same distance the animal naturally maintains during locomotion. In this setup, the spider is unable to move or change orientation, but its intended behavior is transferred to the ball. By recording the latter with a high-speed camera and analyzing the videos with special software (FicTrac, Moore et al., 2014) we were able to infer the animal movement “intentions”.

The styrene ball was placed 20 cm away from the computer monitor (27 inches, 60hz, 1080p), with the spider facing directly towards its center. Stimulus presentation was controlled via a custom script written in Python 3.8 (Van Rossum & Drake, 2009) and the packages *Psychopy* (Peirce et al., 2019), *OpenCV* (Bradski, 2000) and *Numpy* (Oliphant, 2006; van der Walt et al., 2011). After 210 seconds of desensitization, where the monitor showed a blank, white canvas and the spider was free to move, the first stimulus appeared. This consisted of a black dot, 4° in diameter, 30° to the left (or to the right for the other half of the spiders) of the screen’s center. The dot wobbled slightly left-right (or up-down for half of the spiders), at a speed of 30° per second in a span of 7°. After 0.7 seconds, the stimulus disappeared. Then, after a 15 second pause (ITI, inter-trial interval), a second stimulus appeared, identical to the previous one (in size, direction, speed, and position). A total of 10 stimuli were presented to each spider to induce habituation. The 11^th^ stimulus appeared in the same position, had the same speed, and was of the same size, but wobbled up-down (or left-right if the first 10 moved up-down). This stimulus was inserted to induce dis-habituation. Stimuli 12-20 were all identical to stimulus 11, and the 21^st^ changed again, back to what stimulus 1 was. The spiders underwent a total of 45 stimuli presentations with a fixed ITI of 15 seconds, alternating between the two motion directions every 10 stimuli.

When detecting an object with their secondary eyes, jumping spiders perform peculiar pivots (whole-body movements), intended to face the target with their primary eyes (Land, 1985; Zurek et al., 2010). Crucially, the pivots are not produced indiscriminately but rather can be selective (De Agrò et al., 2021). We hypothesize that spiders will lower their response rate across repeated presentations of the same stimulus (will habituate) but will produce responses again when the stimulus changes in one of its characteristics (will dis-habituate). These pivots can be observed in the treadmill setup by analyzing the Z-axis rotations of the sphere.

#### 3.2.3 Day 3 and 5: Personality assay

The spiders underwent the first personality testing. The assay was designed to measure the subject’s boldness. This is derived from behavioral proxies, generally associated with acquiring information about the environment (e.g., moving extensively in an unfamiliar area, which is often equivalent to measuring fear or exploration (Walsh & Cummins, 1976). Each individual was gently released into a novel environment (circle-shaped, 140 mm in diameter, platform) equipped with a shelter (i.e., a piece of cardboard covering one-third of the platform, creating an 8 mm high slit allowing the spider to hide, Fig. 1). The platform was positioned in a plastic container (320 mm x 330 mm x 500 mm) filled with water up to the platform brim to prevent the spider from leaving. The distance between the experimental table and the observer was 30 cm. The platform was cleaned after testing each spider to remove the silk dragline (if any) potentially left by the previously tested spider.

**Figure 1.**
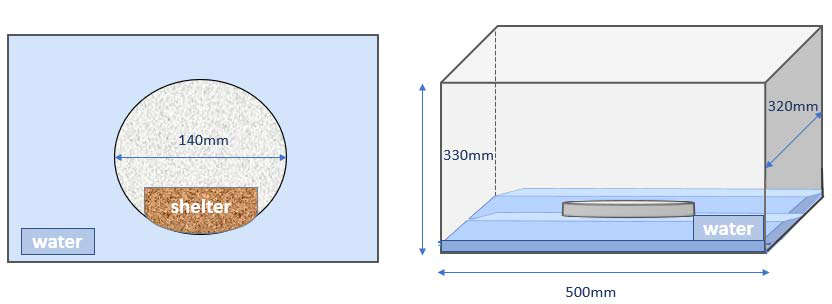
Schematic representation of the arena used in the boldness test. The plastic container is shown from the above (left) and from the side (right). The platform was placed in the center. On top of the platform, a piece of cardboard is placed to provide shelter, indicated by the brown shading.

Each personality assay lasted 20 minutes, which we fully recorded using a Panasonic HC-V180 camera. At the beginning, the spider was given 10 minutes to freely explore the area. Then, the experimenter used long tweezers to “chase” the spider and force it to hide under the shelter. The videos were then analyzed both manually by an experimenter, and automatically with the software YOLACT (Bolya et al., 2022), an image segmentation neural network that we trained to detect the spider across the frames of the videos. We collected the following data: (i) latency to stay under shelter after the disturbance (manually recorded); (ii) presence/absence of jump made towards the water, interpreted as escape attempts (manually recorded); (iii) total walking distance in the first and the last 9 minutes of the test (before and after the disturbance, YOLACT recorded), (iv) average walking speed in the first and the last 9 minutes of the test (before and after the disturbance, YOLACT recorded).

We assumed that bolder individuals would spend less time under the shelter after disturbance, would cover more walking distance in both trial sections, would move at a generally faster speed, and would perform willingness to escape (i.e., jump).

After the first personality assay, the spiders were given a day of rest. The next day the spiders underwent a second personality test identical to the first one, to assess the repeatability of the behavior.

### 3.3 Ethical Note

Each spider used in this study was kept in the laboratory for a total of five days, during which it underwent a visual habituation/dishabituation (H/DH) test and two personality tests (with one-day interval between these personality tests). After spiders were captured, they immediately prepared for the H/DH test by attaching a magnet to their carapace using a UV-activated resin to glue the magnet. Observations show that the magnets did not appear to have a negative effect on the animals. At the end of the test, the magnets were gently removed from the spider’s head. The next day animals were prepared for personality tests. All spiders were released in the same location where they were captured at the end of the study (after the last personality test was completed). The study followed all the legal requirements of the country in which the work was carried out and all institutional guidelines.

### 3.4 Data analysis

The data outputted from FicTrac was extracted using custom software written in Python 3.8, employing the packages *Pandas* (Reback et al., 2020) and *scipy* (Virtanen et al., 2020) From the full array of data, we selected all the frame-wise rotation values around the sphere Z-axis expressed in radians (from now on, ΔZ) in the 200 frames after each stimulus appearance. The camera recorded at 120fps, so 200 frames correspond to 1.66 seconds. Values of ΔZ could be negative or positive, depending on the rotation direction (clockwise or counterclockwise). For the analysis, values representing a rotation congruent with the stimulus position (clockwise for stimuli on the left) were set as positive, and negative otherwise (Note that the ball rotates in the opposite direction to the intended movement of the spider). ΔZs of a spider pivoting towards the stimulus would be characterized by rapid acceleration and then a subsequent deceleration, with the total angle rotated equally to 30° (the stimulus position in respect to the spider facing direction).

The data was then passed on to the statistical software R, version 4.2.2 (R Core Team, 2021), using the library *data.table* (Dowle &d Srinivasan, 2022) given the large dataset. We used the functions from the library *rptR* (Stoffel et al., 2017) to compute the repeatability of four personality measures, taking into account spider’s sex and size.

To test the presence of habituation and dis-habituation, we employed Generalized Additive Models (GAM) from the package *mgcv* (Wood, 2017) with Gaussian error structure. We set ΔZ as the dependent variable; frame from stimulus appearance, stimulus number, and their interaction as smooth terms; whether the stimulus differed from the previous one (dis-habituation stimulus) as a fixed term. We also added subject sex as a fixed term and the interaction between cephalothorax length and width as a smooth term as covariates. Lastly, we included subject identity as a random effect for both intercept and slope. After having observed a steep decrease in response rate after the first 11 stimuli (see results) the data for the subsequent stimuli was discarded and the model was repeated for the first batch only. This was done to exclude the effect of fatigue from the analysis. The measures of personality were modeled as a smooth predictor in the models containing only the first 11 trials and the first 100 frames, given that we observed activity rotation only happening until that point (see results, Fig. 3). We also included the stimulus type (habituation or dis-habituation stimulus) as an interaction variable. We performed a separate GAM for each personality measure. Only the main results are presented here. To see the full analysis, please refer to S1. Raw data is available on S2.

## 4 Results

### 4.1 Personality assay

The following measures were found to be repeatable: the latency to stay under shelter (*R* = 0.316, SE = 0.133, *P* = 0.0169), the presence/absence of a jump in the trial (*R* = 0.30, SE = 0.092, *P* = 0.0134), the total distance walked in the first 9 minutes (*R* = 0.24, SE = 0.122, *P* = 0.0393), the distance walked in the last 9 minutes (*R* = 0.283, SE = 0.115, *P* = 0.0155), and the average speed in the first 9 minutes (*R* = 0.234, SE = 0.089, *P* = 0.0447). On the other hand, the average speed in the last 9 minutes was not repeatable (*R* = 0.175, SE = 0.116, *P* = 0.144). Overall, we observed a high correlation among the behavioral measures of personality. Specifically, the time spent under shelter for both trials correlated negatively with all other measures. The presence/absence of jumps, average speed, and the total distance walked all correlated positively to each-other (Fig. 2).

**Figure 2.**
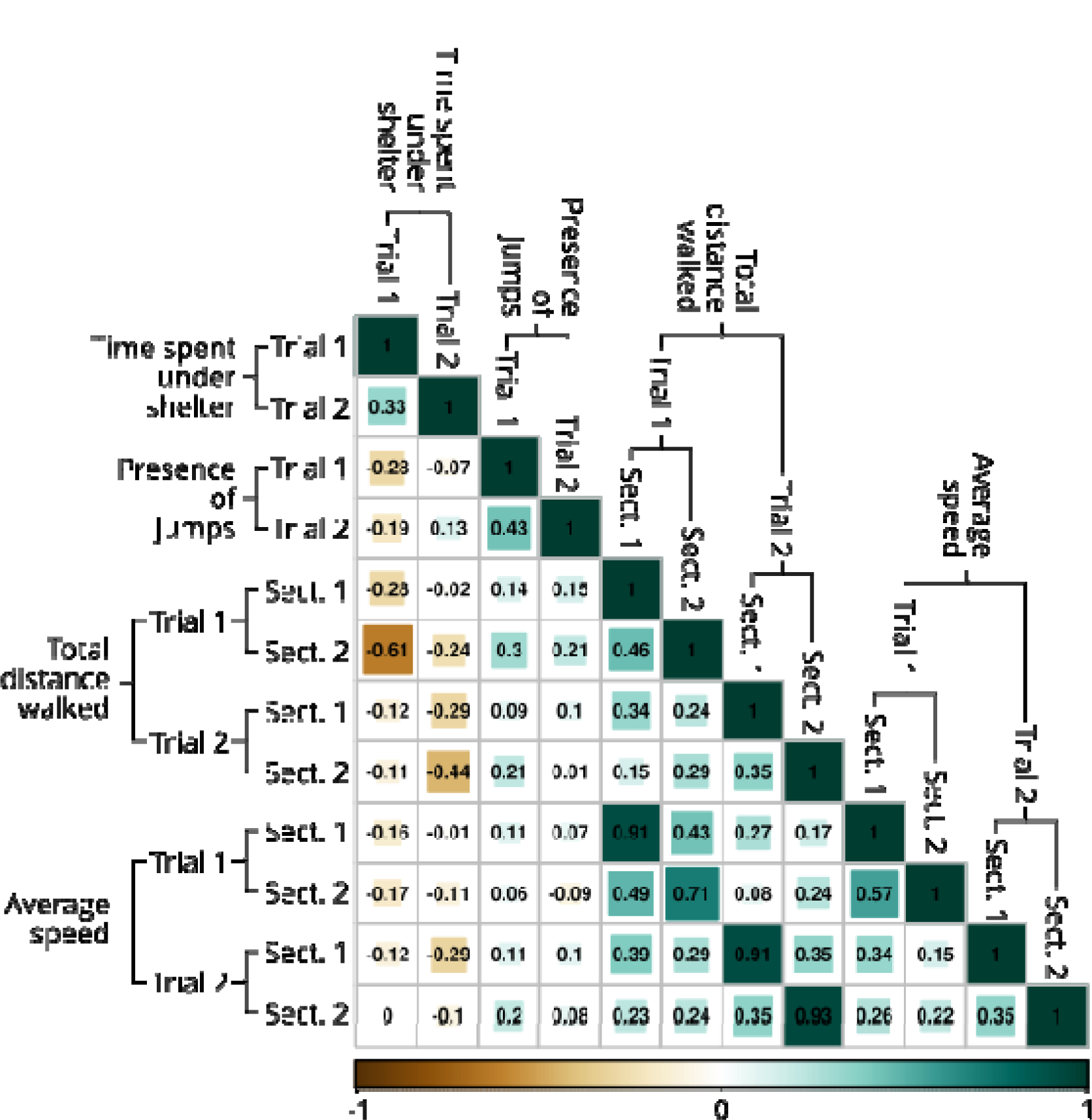
Correlation matrix for all the collected measures of personality. P-values were not computed since the 66 pairwise comparisons would require a multiple-testing correction that is so severe that none of the comparisons would reach statistical significance. However, it is overall appreciable that the latency to stay under shelter is negatively correlated with all other measures, while the latter is positively correlated with each other. Note that for “total distance walked” and “average speed” two measures are available for each trial, corresponding to the first 9 minutes of the test (before the disturbance, Sect. 1) and the last 9 minutes (after the disturbance, Sect. 2).

### 4.2 Habituation/Dis-habituation

There was a significant effect of the interaction between the frame number from stimulus appearance on ΔZ (GAM, EDF = 8.666, RefDF = 8.968, *F* = 85.606, *P* < 0.0001), an effect of stimulus number (EDF = 8.575, RefDF = 8.945, *F* = 22.429, *P* < 0.0001) and of the interaction between the two (GAM, EDF = 14.645, RefDF = 15.713, *F* = 22.789, *P* < 0.0001). The shape of stimulus number (Fig. 3) shows a steep decrease in response (ΔZ) immediately after the first 5 stimuli, remaining around 0 for the rest of the test. However, there are apparent oscillations of increased responses every 11^th^ stimulus, as expected from the habituation test. An increase in the response ruled out fatigue or sensory adaptation. The dis-habituation stimuli (11, 21, 31, 41) did not significantly differ from the others (Estimate = 8.482·10^-5^, SE = 9.854·10^-5^, *t* = 0.861, *P* = 0.389). This was probably due to the immediate drop in response, making later stimuli (>11) all too low to show an appreciable effect.

**Figure 3.**
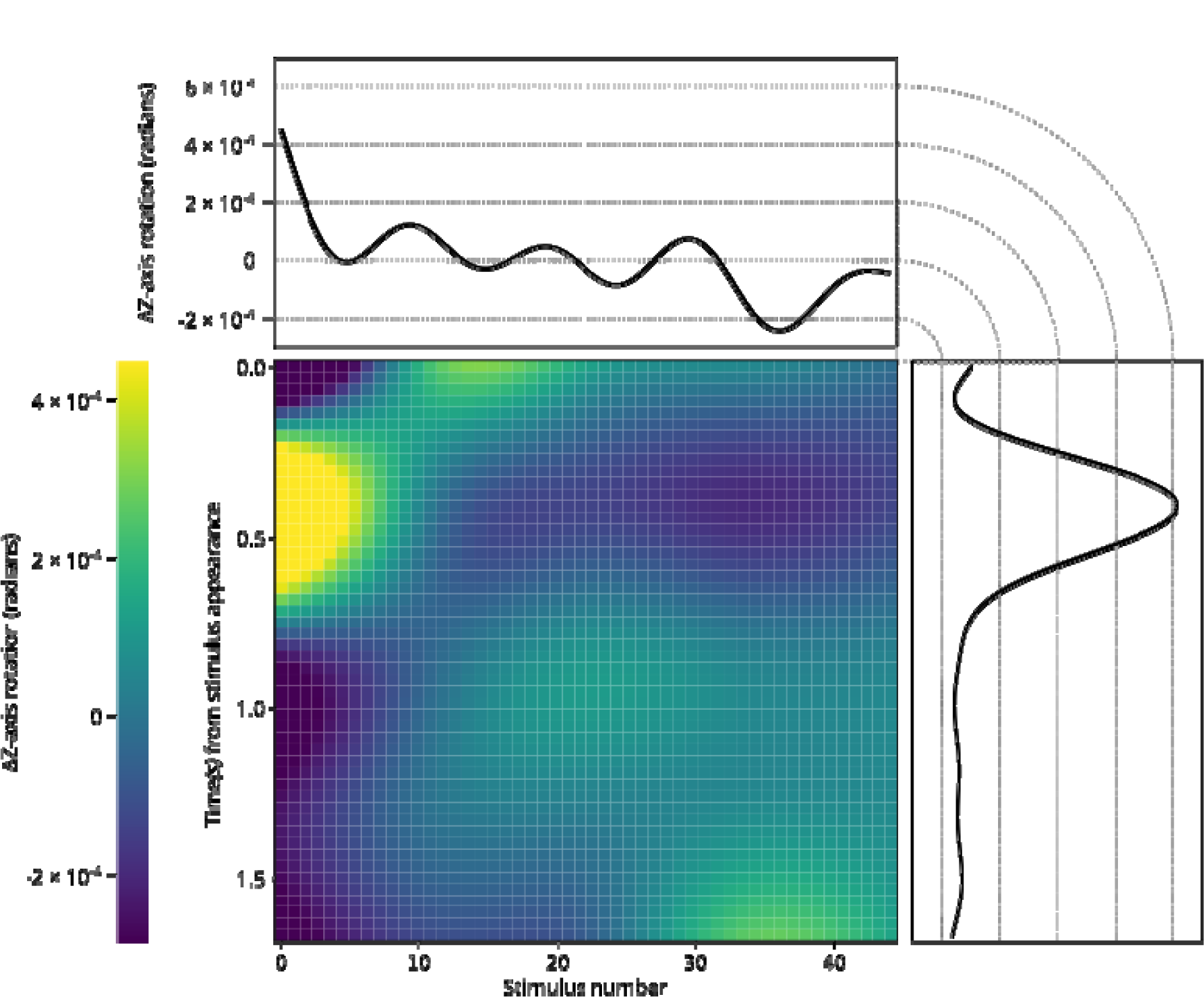
Smooth plots describing the average radians per frame rotational speed of the treadmill, which serves as a proxy for the rotational tendency of the spiders. Positive values indicate rotation towards the position of the stimulus. In the center, a 3D plot describing the interaction between the time from stimulus appearance and the stimulus number. Yellow colour represents the highest observed ΔZ, while dark blue represents the lowest. On the sides are reported the separate curves for clarity. Note that the number and shapes of basic functions generated by the GAM for the interaction plot and the single curves are not the same. The right plot shows rotations as a function of time from stimulus appearance (X-axis). For convenience, time in seconds is displayed on the plot, while the computation was done using the number of frames (120 per second). It is appreciable how after the average stimulus appearance, the spiders rotated towards it, reaching peak speed around 0.5 seconds after appearance and terminating the rotation around 0.25 seconds after that. The top plot shows rotational speed as a function of stimulus number. The first stimulus presents the higher average speed, likely due to a higher number of subjects responding to it. Then, the response rate drops, but a new increase is noticeable every 10 stimuli, when the dis-habituation stimulus is presented.

When restricting the analysis to the first ten stimuli, the interaction between frame number (GAM, EDF = 8.789, RefDF = 8.997, *F* = 83.6, *P* < 0.0001), and stimulus number (EDF = 4.803, RefDF = 5.821, *F* = 12.848, *P* < 0.0001) and their interaction on ΔZ is also significant (GAM, EDF = 13.517, RefDF = 15.189, *F* = 10.736, *P* < 0.0001). Indeed, the response to the dis-habituation stimulus (11) deviates significantly from the expected value (Estimate = 0.0006, SE = 0.0003, *t* = 2.452, *P* = 0.0142).

By looking at the smooth plot of the frame number (Fig. 3), it is possible to see the expected acceleration and deceleration pattern that characterizes the pivot. From around frame 20, the acceleration of the polystyrene ball is appreciable, reaching the maximum speed at around frame 50, and decelerating again until a full stop at around frame 100. Note that this motion is congruent with the stimulus position, as rotation towards it was coded as positive, while rotations in the opposite direction were coded as negative. As such, the random motion would result in an average of 0. Unfortunately, from this analysis, it is impossible to read the more detailed information like the exact percentage of stimuli reacted to, the average angle of rotation, acceleration, or the exact starting point of the rotation. This is because subjects that did not react to a particular stimulus are still included and contribute to the model result as well. In terms of reaction times, the observed start of the acceleration at frame 20 is congruent to what has been described by Zurek and Nelson (2012a), who used a very similar behavioral experiment.

We observed that the time spent under shelter in both trials had a significant effect on ΔZ (GAM, EDF = 1.013, RefDF = 1.013, *F* = 4.589, *P* = 0.0317, Fig. 4). while none of the other measures had a significant effect on ΔZ across the whole experiment, neither on the habituation stimuli only, nor on the dis-habituation ones.

**Figure 4.**
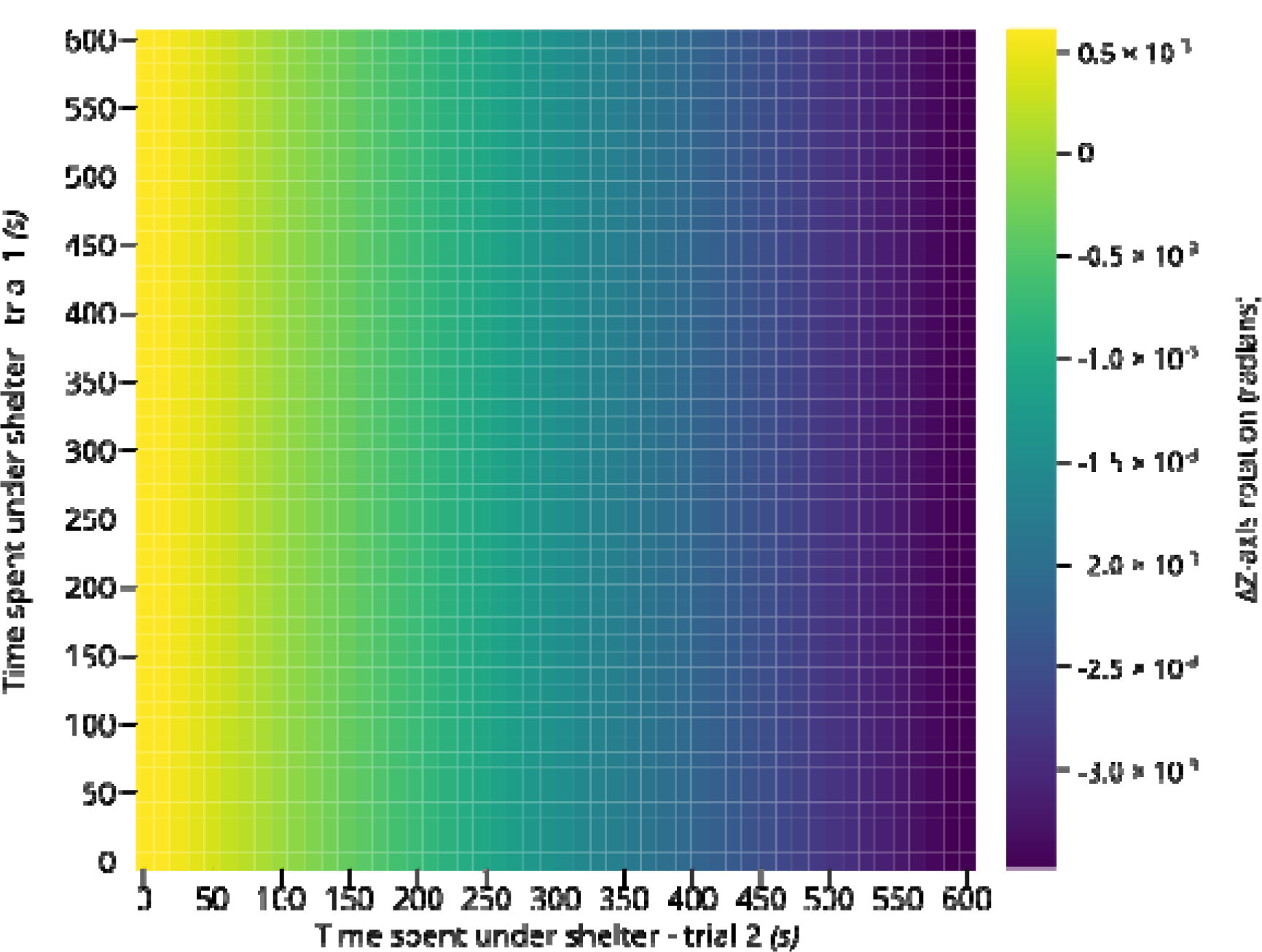
3D smooth plot describing the average radians per frame rotational speed of the treadmill, which serves as a proxy for the rotational tendency of the spiders. Positive values indicate rotation towards the position of the stimulus. The plot describes the effect of the time spent under shelter (taken as a proxy of personality) in both trials on the rotational tendency.

Yellow colour represents the highest observed ΔZ, while dark blue represents the lowest. The plot seems to suggest no relationship between the time spent under shelter in the first trial with the spider performance. On the other hand, a linear relationship is appreciable for the second trial, with an increase in time corresponding with a decrease in rotational tendency.

## 5 Discussion

In this study, we showed that *Menemerus semilimbatus* display a consistent personality trait, specifically boldness, measured patterns using various measures. Furthermore, our finding provides evidence that although the spiders were habituated and dishabituated to visual stimuli, their personality did not play a role in this process.

### 5.1 Consistency in Personality

Our personality test consisted of four different behaviors which, we hypothesized, could be combined into one personality axis, specifically boldness. We hypothesized that these measurements correlate with each other. Specifically, we predicted that bolder individuals would spend less time hiding under shelter, leading to a negative correlation to the presence/absence of jumps toward the water; Additionally, bolder individuals would walk longer distance and exhibit a higher speed of movement.

In accordance with our hypothesis, we found that the latency to stay under shelter after the disturbance in both trials was significantly repeatable, as well as the total walking distance in the first and last 9 minutes, and the average speed during the first 9 minutes. However, we failed to find evidence of significant repeatability in walking speed during the second half of the trials. This was probably due to the fact that the speed calculation for the second half was based on partial data, as, the spiders spent varying amounts of time in shelter after the disturbance, with some individuals never emerging for the remaining 10 minutes of the test. The speed of these spiders was therefore only measured for the time they were outside of the shelter. As spiders alternated between periods of movement and inactivity, calculating speed over short intervals could lead to gross overestimation (if mostly motion sections are observed), or underestimation (if mostly stasis sections are observed). The wide range of potential errors likely prevented us from observing any significant repeatability in this measure.

Jumping behavior can be considered as a willingness to explore based on risk-taking tendencies. Using this assumption, we predicted the occurrence of jump events would correlate positively with other measures. However, there are alternative explanations. For instance, in our experiment, the spider may have jumped toward the water not due to a risk-taking attitude, but as an attempt to escape from the stressful situation. This perspective suggests that shy spiders would be more prone to demonstrate escape (jump). Nevertheless, since we found a positive correlation between the occurrence of jumps and other bold behavioral patterns we measured, it is much more likely that jumping is a component of boldness. Additionally, we observed other spiders inspecting the area before jumping in any direction, suggesting a more risk-prone exploration approach. Therefore, individuals performing the jump are probably not scared but are using this behavior to explore their surroundings.

Unfortunately, there is no universal consensus on which behaviors do and which do not correlate with boldness, making it difficult to define this trait within a broader context. Most behaviors that we recorded in our study could indicate boldness or the opposite, depending on the context in which they are displayed (Mather & Carere, 2019). Locomotion is the flagship example of this uncertainty. In our experiment, we predicted that walking distance and average speed could be direct signs of boldness or related traits. However, some authors found high locomotion was not always an indication of being bolder (Chapman et al., 2011; Coleman & Wilson, 1998; Dellu et al., 1993; Found & St. Clair, 2017; Mazza et al., 2018; Toms et al., 2010). This issue arises from the uncertain definition of boldness as a trait. Some studies suggest that boldness advances locomotor activity (Wilson & Godin, 2010), while others see it as nearly equal to exploratory behavior (Herde & Eccard, 2013) or even co-occurrence of both (Newar & Careau, 2018). Particularly, in our experimental design, we define boldness as the combination of both approaches. This is similar to a behavioral approach and protocol used in a fish model (Mazué et al., 2015). Our results indicate that boldness correlates negatively with latency to stay under shelter (an expression of shyness) and positively with speed and movement patterns.

Despite all the challenges in designing personality measure, the individual differences of these animals appeared stark, especially observing their naive reaction to the trap arena (falcon) during capture where some individuals performed “freezing” behavior without or being less willing to enter the trap while others showed immediate or quick curiosity to explore the trap arena. These inter-individual differences in reactivity level, along with the boldness parameters we measured during the experiment, are most likely a result of a simulated foraging context, although the spiders were not visually fixated on a prey or a predator.

### 5.2 Habituation/Dis-habituation and Personality

The Habituation/Dis-habituation data analysis revealed two main findings. Firstly, we observed a clear decrease in response rate across subsequent stimuli, with a steep drop after just 5 stimuli. Secondly, a key to the current experiment, we observed a significant increase in rotational movement in response to the first dis-habituation stimulus, and a regular pattern in the smooth of stimulus number, with an increase of every 10 stimuli. This demonstrates, as expected, that presenting the animals with a novel stimulus induces an increase in response rate. However, as reported by others (Melrose et al., 2018; Nelson et al., 2019), we also observed a general decrease in response rate, as the response to the dis-habituation stimuli never reached the same magnitude as the response to the very first stimulus. This may be due to the fact that our dis-habituation stimulus becomes the start of a new habituation train, and the spider may maintain a memory of both stimuli after the initial experience, leading to a decreased efficacy in dis-habituating.

We observed the effect of the time spent under shelter on overall activity in the experiment. However, the lack of any other similar effect of any other of the collected measures disqualifies the idea that this is a true effect of personality. Indeed, if it was an effect of personality, it should have been consistent through all, or at least most, boldness measures. Moreover, the effect is on the overall performance, rather than on the habituation or dis-habituation stimuli, and as such linked to the general activity of the animals rather than on their learning processes. In the end, given the small size of the effect and its isolation, we believe this to be a false positive.

Overall, we did not observe any effect of personality in the learning task. We assume that this may be due to the experimental setup we used, where the visual stimuli were formless, leaving the spiders to assign value to them. If the spiders differ in personality, their value attribution process may also vary, resulting in opposite effects for shy and bold individuals. It is possible that bold individuals interpreted the stimuli as a prey and did not habituate because of their boldness and motivation to hunt. On the other hand, shy individuals may have interpreted the stimuli as a predator and did not habituate due to fear and heightened alertness. The response of the two personality types was the same because they interpreted the stimuli differently. Indeed, the link between personality and motivation is related to the context and value (see conceptual papers: Sih & Del Giudice, 2012; Corr et al., 2013 for a review). Alternatively, it is also possible that the spiders did not assign any value to the presented stimuli and viewed them as meaningless moving targets. In this case, the spiders’ reaction to the stimuli would not have been related to their motivation to hunt or escape but rather to their innate tendency to pivot towards moving objects, which is hardwired into the brain and less likely to be affected by personality.

To date, many authors have studied the whole-body saccade behavior of jumping spiders (Bruce et al., 2021; De Agrò et al., 2021; Harland et al., 2012; Jakob et al., 2018; Land, 1972a, 1972b; Zurek et al., 2010; Zurek & Nelson, 2012a, 2012b). Most have reported how the production of pivots is immediate upon stimuli detection. The spiders’ secondary eyes project into dedicated brain areas that occupy most of their visual system (Steinhoff et al., 2020). Given these elements, stimulus detection, recognition, and subsequent turn may be likely foundational to their brain mechanisms, imperturbable by their current motivational state, situation, or indeed personality type as stated before (see above). This remains true also for the dis-habituation stimuli, as motion discrimination seems to be fully functional in the secondary eyes alone (De Agrò et al., 2021), occurring prior to the production of any pivot.

## 6 Conclusion

In summary, our study shows that *Menemerus semilimbatus* spiders display consistent personality traits as evidenced by the behaviors we measured. Bold individuals exhibited longer total distance traveled, faster average speed, less time spent hiding under shelter, and more frequent risk-based behavior (jumps/escape) than shy individuals. These measures were positively correlated, suggesting that they may represent a single behavioral trait-boldness. In addition, we described an effective procedure for visual habituation and dis-habituation in the jumping spider *M. semilimbatus* which allowed us to observe a clear decrease in response rate to stimulus due to habituation as well as a significant increase in rotational movement in the first dis-habituation stimulus.

Lastly, we found no significant correlation between personality and habituation/dis-habituation rates in this spider. In our opinion, this could be due to the fact that bold or shy individuals may have assigned opposite values to ambiguous visual stimuli, interpreting them as either prey or a predator, respectively. This would have placed bold individuals in a hunting context, thereby increasing their attention, while shy individuals would have been in a hiding context, also increasing their attention. Alternatively, the ambiguous stimuli may have had no value at all, and the spiders’ reactions might have relied solely on the hardwired pivot behavior typical to all jumping spiders, making this learning independent of personality. Future research focused on the specific nature of the stimulus would enhance our comparative understanding of how the motivational stage and “pure” response tendency drive an animal’s reaction to such stimuli in relation to the personality axis.

## 7 Author Contributions

N. I. Beydizada, D. Baracchi & M. D. Agrò: Conceptualization; Designing the experiment; Methodology; Data preparation; Writing–Original draft; Writing–Review & editing. N. I. Beydizada & F. Cannone: Material collection & conducting experiments. M. D Agrò & S. Pekár: Data analysis. D. Baracchi, M. D. Agrò & S. Pekár: Supervision. All authors contributed critically to the drafts and gave final approval for publication.

## 8 Data Availability

All data generated during this study are provided as supplementary material (S1 and S2).

## 10 Acknowledgment

N. I. Beydizada was supported by a grant from the Erasmus+ Program (ID: 00216224) for her traineeship in the Cognitive and Behavioural Ecology Lab (CBE), at Florence University, Italy.

